# Ocular and uteroplacental pathology in macaque congenital Zika virus infection

**DOI:** 10.1101/195701

**Authors:** Emma L. Mohr, Lindsey N. Block, Christina M. Newman, Laurel M. Stewart, Michelle Koenig, Matthew Semler, Meghan E. Breitbach, Leandro B.C. Teixeira, Xiankun Zeng, Andrea M. Weiler, Gabrielle L. Barry, Troy H. Thoong, Gregory J. Wiepz, Dawn M. Dudley, Heather A. Simmons, Andres Mejia, Terry K. Morgan, M. Shahriar Salamat, Sarah Kohn, Kathleen M. Antony, Matthew T. Aliota, Mariel S. Mohns, Jennifer M. Hayes, Nancy Schultz-Darken, Michele L. Schotzko, Eric Peterson, Saverio Capuano, Jorge E. Osorio, Shelby L. O’Connor, Thomas C. Friedrich, David H. O’Connor, Thaddeus G. Golos

**Affiliations:** Department of Pediatrics, University of Wisconsin-Madison; Department of Pathology and Laboratory Medicine, University of Wisconsin-Madison; Department of Pathobiological Sciences, University of Wisconsin-Madison; United States Army Medical Research Institute of Infectious Diseases, Fort Detrick, Frederick, Maryland; Wisconsin National Primate Research Center, University of Wisconsin-Madison; Departments of Pathology and Obstetrics & Gynecology, Oregon Health & Science University; Department of Obstetrics and Gynecology, University of Wisconsin-Madison; Department of Comparative Biosciences, University of Wisconsin-Madison; Department of Radiology, University of Wisconsin-Madison

**Author notes:** These authors contributed equally to this work (Emma L. Mohr, Lindsey N. Block, Christina M. Newman). Correspondence and request for materials should be addressed to E.L.M. or T.G.G.

## Abstract

Congenital Zika virus (ZIKV) infection impacts fetal development and pregnancy outcomes. We infected a pregnant rhesus macaque with a Puerto Rican ZIKV isolate in the first trimester. The pregnancy was complicated by preterm premature rupture of membranes (PPROM) and fetal demise 49 days post infection (gestational day 95). Significant pathology at the maternal-fetal interface included acute chorioamnionitis, placental infarcts, and leukocytoclastic vasculitis of the myometrial radial arteries. ZIKV RNA was disseminated throughout the fetus tissues and maternal immune system at necropsy, as assessed by quantitative RT-PCR for viral RNA. Replicating ZIKV was identified in fetal tissues, maternal lymph node, and maternal spleen by fluorescent in situ hybridization for viral replication intermediates. Fetal ocular pathology included a choroidal coloboma, suspected anterior segment dysgenesis, and a dysplastic retina. This is the first report of ocular pathology and prolonged viral replication in both maternal and fetal tissues following congenital ZIKV infection in rhesus macaques. PPROM followed by fetal demise and severe pathology of the visual system have not been described in macaque congenital infection previously; further nonhuman primate studies are needed to determine if an increased risk for PPROM is associated with congenital Zika virus infection.

**Author summary:** A ZIKV infection during pregnancy is associated with malformations in fetal development including, but not limited to, ocular and brain anomalies, such as microcephaly, and stillbirth. The development of an accurate pregnancy model to study the effects of ZIKV will provide insight into vertical transmission, ZIKV tissue distribution, and fetal injury and malformations. Non-human primates closely resemble human in terms of the reproductive system, immunity, placentation and pregnancy. Our study demonstrates that the rhesus macaque is a compelling model in which to study ZIKV during pregnancy due to similar outcomes between the human and rhesus macaque. These similarities include prolonged viremia, vertical transmission, adverse pregnancy outcomes and fetal pathology, including defects in the visual system.

## Introduction

First isolated from a febrile rhesus macaque in Uganda in 1947, Zika virus (ZIKV) generally did not result in recognized widespread clinical disease in subsequent outbreaks across Asia and the South Pacific, until late 2015, when clinicians in Northeast Brazil reported a surge in babies born with severe birth defects (1). By early 2016, the US Centers for Disease Control and Prevention (CDC) asserted that there was a causal relationship between prenatal ZIKV infection and serious brain anomalies including microcephaly (2). The constellation of fetal and neonatal abnormalities and birth defects associated with ZIKV infection *in utero* is designated congenital Zika syndrome (CZS) (3-9). Characteristics of CZS include ocular anomalies, brain anomalies, stillbirth, cranial dysmorphologies, musculoskeletal contractures and neurologic sequelae (10). Infection during the first trimester increases the risk for birth defects (5) because critical cell proliferation and differentiation occurs during this trimester (11). One striking characteristic of CZS is a high frequency of ocular malformations, observed in as many as 55% of infants with evidence of congenital ZIKV infection and microcephaly (12, 13). Multiple case reports and case series have identified infants with ocular anomalies, which include macular pigment mottling, optic nerve hypoplasia, chorioretinal and iris coloboma, lens subluxation, retinal vascular abnormalities, cataracts and maculopathy (5, 14-21). Specific retinal defects include retinal thinning, discontinuity of the retinal pigment epithelium, and colobomatous-like excavation in the neurosensory retina, retinal pigment epithelium and choroid in multiple infants (17). Because the retina develops as an outpocketing from the neural tube (22), the presence of retinal lesions implies CNS damage even without brain abnormalities.

Other recognized outcomes of congenital ZIKV infection are miscarriage, stillbirth and PPROM (23-26). The etiology of PPROM is multifactorial (27). Prenatal ZIKV infection in the first trimester of gestation results in up to 25% of pregnancies with miscarriage, fetal loss or stillbirth, with lower frequencies in the second and third trimesters in a study including 125 pregnancies (28). The CDC reports 15 fetal demise cases with birth defects out of 4,695 live births in women with confirmed ZIKV infection (29). However, this number likely does not capture the total number of fetal demises following congenital ZIKV infection because it does not include fetuses without overt birth defects even though there may be vertical transmission, or early pregnancy losses from women who were not aware of infection, or never sought a diagnosis. The pathophysiology of preterm birth or fetal loss before viability following congenital ZIKV infection has not been defined. In murine models, pregnancy following a systemic viral infection can result in an ascending bacterial uterine infection, inflammation, and preterm birth (30). It has also been reported that viral persistence of ZIKV in the lower female genital tract in the rhesus monkey is prolonged in animals treated with Depo-Provera, a synthetic progestogen (31). The specific etiology of adverse pregnancy outcomes in congenital ZIKV infection, however, is yet to be defined and requires further study.

One novel feature of ZIKV infection is the persistence of both ZIKV RNA (32-37) and replication competent virus in body fluids (33, 38) for weeks after infection. ZIKV RNA has been identified in semen between 3-188 days after infection (39, 40) with a median of 34 days (32), in urine up to 29 days after infection (41) with a median of 8 days (32), in saliva up to 29 days after infection (41), and in serum with a median of 14 days (32). Both ZIKV RNA and infectious particles have been isolated from breast milk 2 days after infection (42). In comparison, RNA from dengue virus (DENV), the flavivirus most closely related to ZIKV, has only been isolated from urine up to 3-4 weeks after infection (43); no DENV RNA has been isolated from semen, prolonged plasma viremia has only been reported in hematopoietic stem cell recipients (44, 45), and only DENV RNA has been isolated from breast milk around the time of acute infection (46). Defining the body fluid and tissue persistence of ZIKV is critical to the development of public health recommendations and solid organ and hematopoietic stem cell transplant guidelines. Non-human primate (NHP) models have begun to define the tissue distribution of ZIKV following infection because defining tissue distribution in humans is not possible. Following ZIKV infection in nonpregnant NHPs, ZIKV has been identified in multiple tissues up to 35 days after infection, including the brain, spinal cord, eye, spleen, lymph nodes, muscles and joints (47, 48) and in cerebrospinal fluid (CSF) up to 42 days after infection (48), suggesting that one of these tissues may support prolonged ZIKV replication. Since prolonged ZIKV viremia is a feature of ZIKV infection during pregnancy, we hypothesized that ZIKV tissue persistence would be longer in pregnant NHPs compared to nonpregnant NHPs. Indeed, following ZIKV infection in pregnant NHPs, ZIKV RNA detection in plasma is prolonged (47) and can be detected up to 70 days after infection (49), far longer than the plasma viremia duration reported for nonpregnant NHPs (48, 50).

NHP models of both congenital infection and tissue distribution following ZIKV infection provide insight into the pathophysiology of ZIKV infection not possible through epidemiological and clinical human studies. As with humans, the rhesus macaque placenta has a hemochorial placentation with extensive endovascular invasion of the maternal endometrial spiral arterioles and arteries and innate immune cellular populations homologous with that found in the human decidua (51-53). There are multiple similarities between human and NHP ZIKV infection natural history, including the duration of viremia and viruria (47, 48, 50, 54), robust neutralizing antibody responses (47, 50, 54, 55), vertical transmission (49), and fetal pathology (49, 56). To define the tissue distribution of ZIKV and fetal pathology following infection with a clinically relevant Puerto Rican isolate of ZIKV, we infected a pregnant rhesus macaque in the first trimester and performed a necropsy of the dam and fetus to comprehensively define maternal and fetal viral tissue distribution following spontaneous fetal death 49 days post-infection. Here, we describe the pregnancy outcome, maternal and fetal viral tissue distribution, and fetal pathology associated with first trimester ZIKV infection in a case of fetal demise.

## Materials & Methods

### Study Design

A 3.8 year old, primigravida rhesus macaque (*Macaca mulatta*) of Indian ancestry was infected subcutaneously with 1x10^4^ PFU Zika virus/H.sapiens-tc/PUR/2015/PRVABC59_v3c2 (PRVABC59) during the first trimester, 46 days gestation (term 165±10 days). This macaque was part of the Specific Pathogen Free (SPF) colony at the Wisconsin National Primate Research Center (WNPRC) and was free of Macacine herpesvirus 1 (Herpes B), Simian Retrovirus Type D (SRV), Simian T-lymphotropic virus Type 1 (STLV), and Simian Immunodeficiency Virus (SIV).

### Ethics

All monkeys are cared for by the staff at the WNPRC in accordance with the regulations and guidelines outlined in the Animal Welfare Act and the Guide for the Care and Use of Laboratory Animals and the recommendations of the Weatherall report (https://royalsociety.org/topics-policy/publications/2006/weatherall-report/). This study was approved by the University of Wisconsin-Madison Graduate School Institutional Animal Care and Use Committee (animal protocol number G005401).

### Care & Use of Macaques

The female monkey described in this report was co-housed with a compatible male and observed daily for menses and breeding. Pregnancy was detected by ultrasound examination of the uterus at approximately 20-24 gestation days (gd) following the predicted day of ovulation. The gd was estimated (+/- 2 days) based on the dam’s menstrual cycle, observation of copulation, and the greatest length of the fetus at initial ultrasound examination which was compared to normative growth data in this species (57). For physical examinations, virus inoculations, some ultrasound examinations, blood and swab collections, the dam was anesthetized with an intramuscular dose of ketamine (10 mg/kg). Blood samples from the femoral or saphenous vein were obtained using a vacutainer system or needle and syringe. The pregnant macaque was monitored daily prior to and after inoculation for any clinical signs of infection (e.g., diarrhea, inappetence, inactivity and atypical behaviors). This macaque developed chronic diarrhea prior to conception and was treated daily with oral tylosin throughout pregnancy.

### Inoculation and monitoring

ZIKV strain PRVABC59 (GenBank: KU501215), originally isolated from a traveler to Puerto Rico and passaged three times on Vero cells (American Type Culture Collection (ATCC): CCL-81), was obtained from Brandy Russell (CDC, Ft. Collins, CO). Virus stocks were prepared by inoculation onto a confluent monolayer of C6/36 cells (*Aedes albopictus* larval cells; ATCC: CCL-1660) with two rounds of amplification. The inoculating stock was prepared and validated as previously described (49, 50). The animal was anesthetized as described above, and 1 mL of inoculum at 1 x 10^4^ PFU dilution in PBS was administered subcutaneously over the cranial dorsum. Post-inoculation, the animal was closely monitored by veterinary and animal care staff for adverse reactions or any signs of disease.

### Pregnancy monitoring and fetal measurements

Weekly ultrasounds were conducted to observe the health of the fetus and to obtain measurements including fetal femur length (FL), biparietal diameter (BPD), head circumference (HC), and heart rate, with methods as previously described (49). Growth curves were developed for FL, BPD, and HC (58). Mean measurements and standard deviations at specified days of gestation in rhesus macaques were retrieved from Tarantal et al. (57) and the data were plotted against normative data for fetal rhesus macaques (58). The actual growth measurements were obtained from weekly ultrasound data and used to retrieve the predicted growth measurement by plotting the obtained experimental growth measurement against the growth curves. Data were then graphed as actual gestation age versus predicted gestation age to depict rate of growth compared to uninfected, control rhesus macaques (method described previously in (49)). Doppler ultrasounds to measure fetal heart rate were performed as requested by veterinary staff.

### Amniocentesis

For the amniocentesis procedures reported, animals were shaved, and the skin was prepped with Betadyne^®^ solution, and sterile syringes, needles and gloves were used during the amniocentesis procedure. Under real-time ultrasound guidance, a 22 gauge, 3.5 inch Quincke spinal needle was inserted into the amniotic sac as described previously (49). The first 1.5-2 mL of fluid was discarded due to potential maternal contamination, and an additional 3-4 mL of amniotic fluid was collected in a new sterile syringe for viral qRT-PCR analysis as described elsewhere (50). These samples were obtained at the gestational ages specified in Figure 1. All fluids were free of any blood contamination.

**Figure 1.**
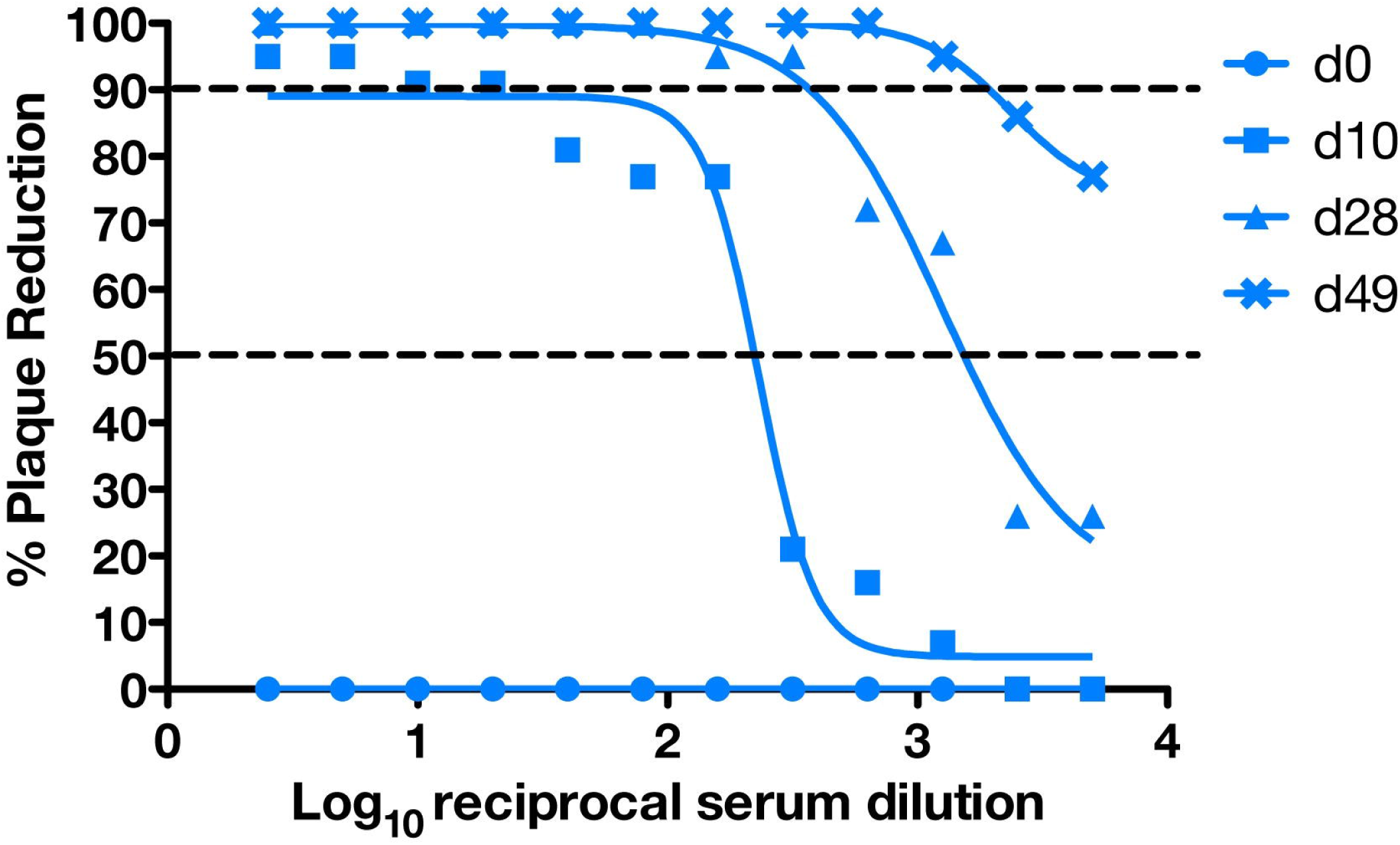
Timeline depicting body fluid sampling and procedures throughout pregnancy. Blood, urine, saliva, amniotic fluid, and CSF were collected as indicated in the schedule above, and ultrasounds were performed weekly. The axes are not drawn to scale.

### vRNA isolation from body fluids and tissues

RNA was isolated from maternal plasma, urine, saliva and amniotic fluid using the Viral Total Nucleic Acid Purification Kit (Promega, Madison, WI, USA) and from maternal and fetal tissues using the Maxwell 16 LEV simplyRNA Tissue Kit (Promega, Madison, WI) on a Maxwell 16 MDx instrument as previously reported (50). Fetal and maternal tissues were processed with RNAlater^®^ (Invitrogen, Carlsbad, CA) according to manufacturer protocols. 20-40 mg of each tissue was homogenized using homogenization buffer from the Maxwell 16 LEV simplyRNA Tissue Kit and two 5 mm stainless steel beads (Qiagen, Hilden, Germany) in a 2 mL snap-cap tube, shaking twice for 3 minutes at 20 Hz each side in a TissueLyser (Qiagen, Hilden, Germany). The isolation was continued according to the Maxwell 16 LEV simplyRNA Tissue Kit protocol, and samples were eluted into 50 μL RNase free water.

### Viral quantification by plaque assay

Titrations for replication competent virus quantification of amniotic fluid was completed by plaque assay on Vero cell cultures as described previously (50). Vero cells were obtained from American Type Culture Collection (CCL-81), were not further authenticated and were not specifically tested for mycoplasma. Duplicate wells were infected with 0.1ml of aliquots from serial 10-fold dilutions in growth media and virus was adsorbed for 1 h. Following incubation, the inoculum was removed, and monolayers were overlaid with 3 ml containing a 1:1 mixture of 1.2% oxoid agar and 2 DMEM (Gibco, Carlsbad, CA, USA) with 10% (vol/vol) FBS and 2% (vol/vol) penicillin/streptomycin. Cells were incubated at 37°C in 5% CO_2_ for 4 days for plaque development. Cell monolayers then were stained with 3 ml of overlay containing a 1:1 mixture of 1.2% oxoid agar and 2 DMEM with 2% (vol/vol) FBS, 2% (vol/vol) penicillin/streptomycin and 0.33% neutral red (Gibco). Cells were incubated overnight at 37°C and plaques were counted.

### Plaque Reduction Neutralization test (PRNT)

Macaque serum samples were screened for ZIKV neutralizing antibodies utilizing a plaque reduction neutralization test (PRNT). End point titrations of reactive sera, utilizing a 90% cutoff (PRNT90), were performed as described (59) against ZIKV strain PRVABC59. Briefly, ZIKV was mixed with serial 2-fold dilutions of serum for 1 hour at 37°C prior to being added to Vero cells and neutralization curves were generated using GraphPad Prism software (La Jolla, CA). The resulting data were analyzed by nonlinear regression to estimate the dilution of serum required to inhibit both 90% and 50% of infection.

### Maternal and neonatal necropsy

At 49 days post infection (dpi) (gd 95), no fetal heartbeat was detected. The dam was sedated, euthanized, and sterile instruments were used for the dissection and collection of all maternal, fetal, and maternal-fetal interface tissues during the gross post-mortem examination. Amniotic fluid was aspirated with a syringe and needle inserted through the uterine wall into the lumen. Each tissue was collected with a unique set of sterile instruments and placed in a separate sterile petri dish before transfer to appropriate containers for viral RNA analysis and histology, to prevent cross-contamination between tissues. Tissue distribution for subsequent analysis was as previously described (49).

### Histology

For general pathology, tissues were fixed in 4% PFA as for IHC, routinely processed and embedded in paraffin. Paraffin sections (5 μm) were stained with hematoxylin and eosin (H&E). Two veterinary pathologists were blinded to vRNA findings when tissue sections were evaluated microscopically. Lesions in each tissue were described and assigned morphologic diagnoses as described previously (49). Photomicrographs were obtained using brightfield microscopes Olympus BX43 and Olympus BX46 (Olympus Inc., Center Valley, PA) with attached Olympus DP72 digital camera (Olympus Inc.) and Spot Flex 152 64 Mp camera (Spot Imaging, Sterling Heights, MI), and captured using commercially available image-analysis software (cellSens DimensionR, Olympus Inc. and Spot software 5.3). Uteroplacental pathology was specifically performed by an experienced placental pathologist (T.K.M.).

### In situ hybridization

In situ hybridization (ISH) was conducted with tissues fixed in 4% PFA, and alcohol processed and paraffin embedded, as for IHC. ISH probes against Zika genome were purchased commercially (Advanced Cell Diagnostics, Cat No. 468361, Newark, California, USA). ISH was performed using the RNAscope^®^ Red 2.5 Kit (Advanced Cell Diagnostics, Cat No. 322350) according to the manufacturer’s instructions. Briefly, after deparaffinization with xylene, a series of ethanol washes, and peroxidase blocking, sections were heated in boiling antigen retrieval buffer for 15 minutes and then digested by proteinase K (2.5 ug/ml, to completely cover the section) for 16 minutes at 40°C. Sections were exposed to ISH target probe and incubated at 40°C in a hybridization oven for 2 h. After rinsing, ISH signal was amplified using company-provided Preamplifier and Amplifier conjugated to horseradish peroxidase (HRP), and incubated with a red substrate-chromogen solution for 10 min at room temperature.

### Multiplex fluorescent in situ hybridization

Multiplex fluorescent in situ hybridization (mFISH) was conducted with tissues fixed in 4% PFA as for IHC. mFISH was performed using the RNAscope^®^ Fluorescent Multiplex Kit (Catalog # 320850, Advanced Cell Diagnostics) according to the manufacturer’s instructions with modifications. Probes with C1 channel (Cat# 468361, red) targeting ZIKV positive sense RNA and probes with C3 channel (Cat# 467911, green) targeting ZIKV negative sense RNA were synthesized by Advanced Cell Diagnostics. Paraformaldehyde fixed paraffin embedded rhesus monkey fetus tissue sections underwent deparaffinization with xylene and a series of ethanol washes. These tissue sections were treated with 0.1% Sudan Black B (Sigma-Aldrich, St. Louis, MO, USA) to reduce autofluorescence, heated in antigen retrieval buffer (Citrate buffer with pH 6.0), and digested by proteinase. Sections were exposed to ISH target probes and incubated at 40°C in a hybridization oven for 2 h. After rinsing, ISH signal was amplified using company-provided Pre-amplifier and Amplifier conjugated to fluorescent dye. Sections were counterstained with 4’, 6-diamidino-2-phenylindole (DAPI, Thermo Fisher Scientific, Waltham, MA, USA), mounted, and stored at 4°C until image analysis. mFISH images were captured on an LSM 880 Confocal Microscope with Airyscan (Zeiss, Oberkochen, Germany) and processed using open-source ImageJ software (National Institutes of Health, Bethesda, MD, USA).

### Placental alpha microglobulin-1 (PAMG-1) immunochromatographic assay

A PAMG-1 immunochromatographic assay (AmniSure^®^ ROM (Rupture of [fetal] Membranes) test, Qiagen, Boston, MA, FMRT-1-10-US) was performed according to the manufacturer’s protocol with urine and amniotic fluid samples that had been stored at -80°C. A sterile polyester swab, provided by the manufacturer, was inserted into a tube containing the sample fluid for 1 minute. The swab was then added to the solvent microfuge tube and rotated by hand for 1 minute. Finally, the test strip was placed into the solvent and incubated at room temperature for 10 minutes before the test strip was read and photographs were taken. A term amniotic fluid sample was the positive control and non-pregnant urine was the negative control. Open-source ImageJ software was used to measure the relative pixel density of each band (control and test band) (National Institutes of Health, Bethesda, MD, USA). The pixel density of each band was measured, the background density was subtracted, and the relative pixel density of each test band was calculated by subtracting the control band density from the test band density.

### Insulin-like growth factor-binding protein 1 (IGFBP-1) ELISA

An IGFBP-1 ELISA kit (Abcam, Cambridge, MA, ab100539) was used to determine if a marker for amniotic fluid was detectable in maternal urine. The protocol was followed as specified by the manufacturer and all samples were frozen undiluted at -80°C until use. Duplicates were run for the standards, samples, positive, and negative controls. A term amniotic fluid sample was used as the positive control and male urine and non-pregnant female urine were used as negative controls. All urines were diluted 1:5000 and all amniotic fluid samples were diluted 1:20,000. Immediately upon addition of the stop solution the plate was read at 450 nm. A standard curve was calculated from the average of each standard. This standard curve equation was used to calculate the concentration of each sample.

### Data availability

Primary data that support the findings of this study are available at the Zika Open-Research Portal (https://zika.labkey.com/project/OConnor/ZIKV-019/begin.view?). Zika virus/H.sapiens-tc/PUR/2015/PRVABC59-v3c2 sequence data have been deposited in the Sequence Read Archive (SRA) with accession code SRX2975259. The authors declare that all other data supporting the findings of this study are available within the article and its supplementary information files.

## Results

### Pregnancy outcome

A pregnant rhesus macaque was subcutaneously inoculated with 1x10^4^ PFU ZIKV-Puerto Rico at gd46. She had no fever, rash, or inappetence detected following inoculation. The pregnancy was monitored by ultrasonography, vRNA titers in blood, and urine samples, and neutralizing antibody titers at multiple times throughout pregnancy; amniotic fluid and maternal CSF were collected at several time points (Figure 1).

Maternal plasma viremia was detected from days 1 through 18 post-infection, peaking at 5 dpi with 2.55 x 10^5^ vRNA copies per mL, and was also detected at 24 dpi (Figure 2). At 21 dpi the viral load dropped below 100 vRNA copies per mL, the limit of quantification of the qRT-PCR assay. Amniotic fluid at 28 dpi had a viral load of 1.84 x 10^4^ vRNA copies per mL. The amniotic fluid was reported as clear, and a plaque assay performed on the amniotic fluid was negative (data not shown). Saliva samples remained negative throughout pregnancy (data not shown). CSF samples taken at 7, 14, and 49 dpi were all negative. ZIKV RNA was first detected in a passively collected urine sample (i.e. in pan at the bottom of the cage) at 42 dpi, with a concentration of 7 x 10^4^ vRNA copies per mL, and was present in the urine until euthanasia at 49 dpi.

**Figure 2.**
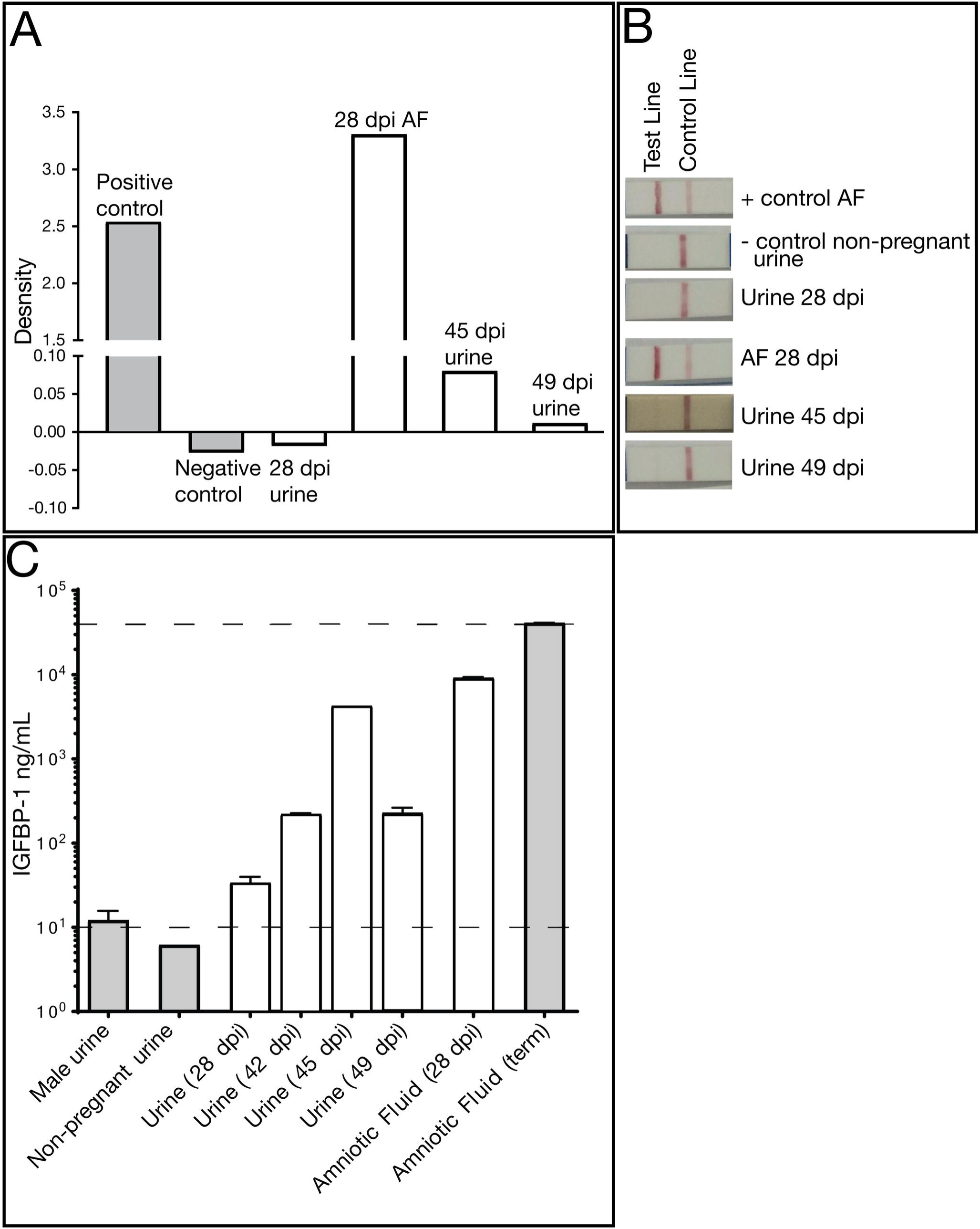
ZIKV vRNA levels in maternal body fluids. vRNA was measured by quantitative RT-PCR in plasma, urine, amniotic fluid and CSF. The limit of assay quantification is 100 copies/mL and the limit of detection is 33 copies/mL.

In addition to vRNA in body fluids, the development of maternal ZIKV-specific antibodies was assessed. Plaque reduction neutralization tests (PRNT) were performed on serum collected at 10, 28, and 49 dpi. All post-infection time points demonstrated the presence of ZIKV-specific neutralizing antibodies with an increasing concentration of neutralizing antibodies throughout the post-infection period (Figure 3).

**Figure 3.**
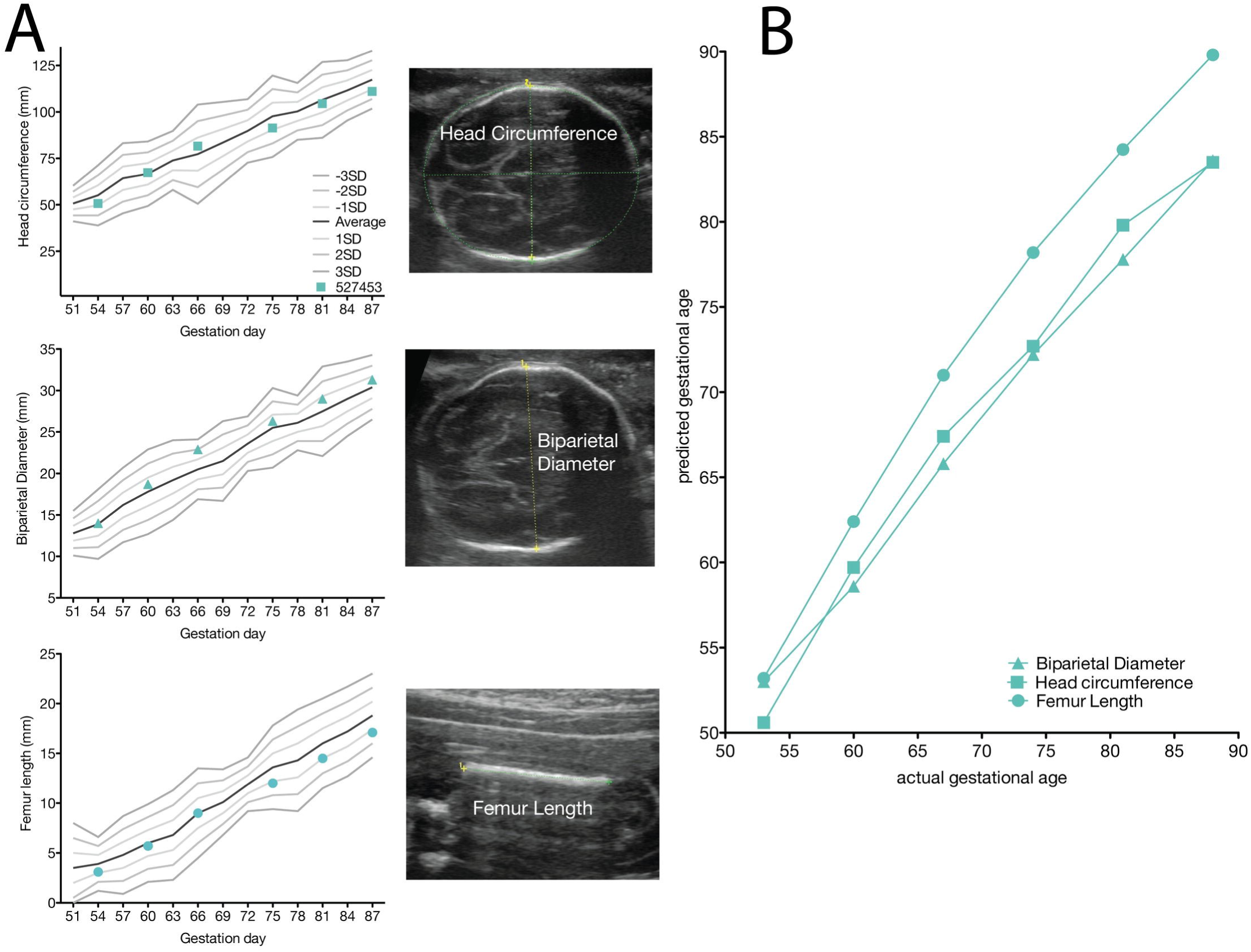
Neutralizing antibody titers following ZIKV infection. PRNT titers were measured pre and post infection. The x-axis represents the reciprocal serum dilution (log_10_) and the y-axis represents the percent reduction. The dashed lines indicate 90% and 50% reduction.

Because identifying urine vRNA so long after infection was unexpected, and its presence coincided with the presence of vRNA in the amniotic fluid, we wanted to determine whether the passively collected urine contained amniotic fluid, a potential harbinger of an adverse pregnancy outcome. We performed an AmniSure^®^ test, which detects an amniotic fluid protein, placental alpha microglobulin-1, (PAMG-1), and determined that urine contained detectable PAMG-1 (Figure 4). As expected, the 28 dpi amniotic fluid was positive for PAMG-1, as was the positive control term amniotic fluid from a different animal. The 28 dpi urine (collected just prior to the amniocentesis) was negative for PAMG-1, however the 45 and 49 dpi urine samples were positive for PAMG-1. The negative control was a non-pregnant urine sample. AmniSure^®^ is not a quantitative test but the result suggested there was amniotic fluid in the urine samples at 45 and 49 dpi. To confirm this finding, we performed an insulin-like growth factor binding protein-1 (IGFBP-1) ELISA on the animal’s pan-collected urine and amniotic fluid samples, along with appropriate controls. IGFBP-1 is a 25 kB protein synthesized and secreted by the fetal liver and maternal decidua, and is present in amniotic fluid from the second trimester of pregnancy until full term (60). It is not found in urine. In the pregnant animal, IGFBP-1 was detected in pan-collected urine at levels similar to that in amniotic fluid alone, confirming the presence of amniotic fluid-specific protein in the urine (Figure 4). The IGFBP-1 levels in urine from this dam were higher than negative control urine samples (urine from a male and a nonpregnant female) but lower than amniotic fluid from a control macaque late in gestation, which is consistent with dilution from urine from passive collection. Thus, the presence of amniotic fluid in the urine is consistent with premature rupture of membranes.

**Figure 4.**
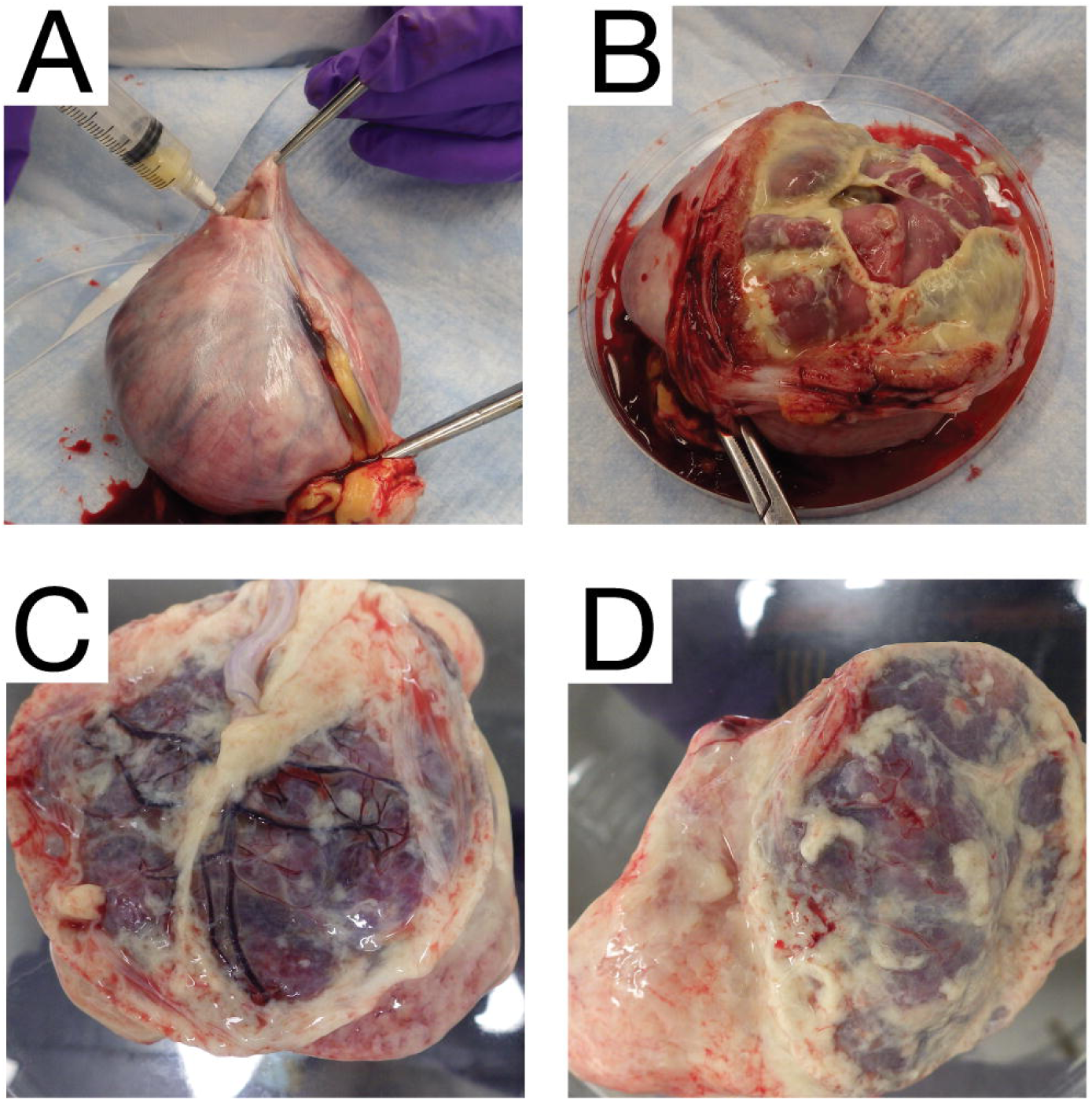
Amniotic fluid (AF) markers confirm rupture of membranes. (A) An AmniSure^®^ test, which measures PAMG-1 protein, was performed on pan urine collection (28 dpi, 45 dpi, 49 dpi) and AF (28 dpi) from the pregnant animal. Nonpregnant control animal urine and pregnant animal AF are included as controls. (B) Relative pixel density of the Amnisure^®^ test strip test band and control band. (C) Amniotic fluid protein IGFBP-1 ELISA. Body fluids from the pregnant animal (pan urine collection 28, 42, 45, 49 dpi and AF 28 dpi), nonpregnant negative control male and female urine samples, amniotic fluid from a control pregnancy were evaluated for the presence of IGFBP-1. In Panels B and C, white bars denote body fluids from the experimental animal and grey bars denote control fluids from other animals in the colony.

### Ultrasonography

The fetus displayed typical growth in all parameters when compared with normative data (Figure 5) (58). Plotting the predicted gestational age (pGA) vs. the clinically estimated (actual) gestational age (aGA) can reveal changes in the trajectory of a specific growth parameter (49, 57); this analysis did not reveal any growth trajectory anomalies (Figure 5B growth chart).

**Figure 5.**
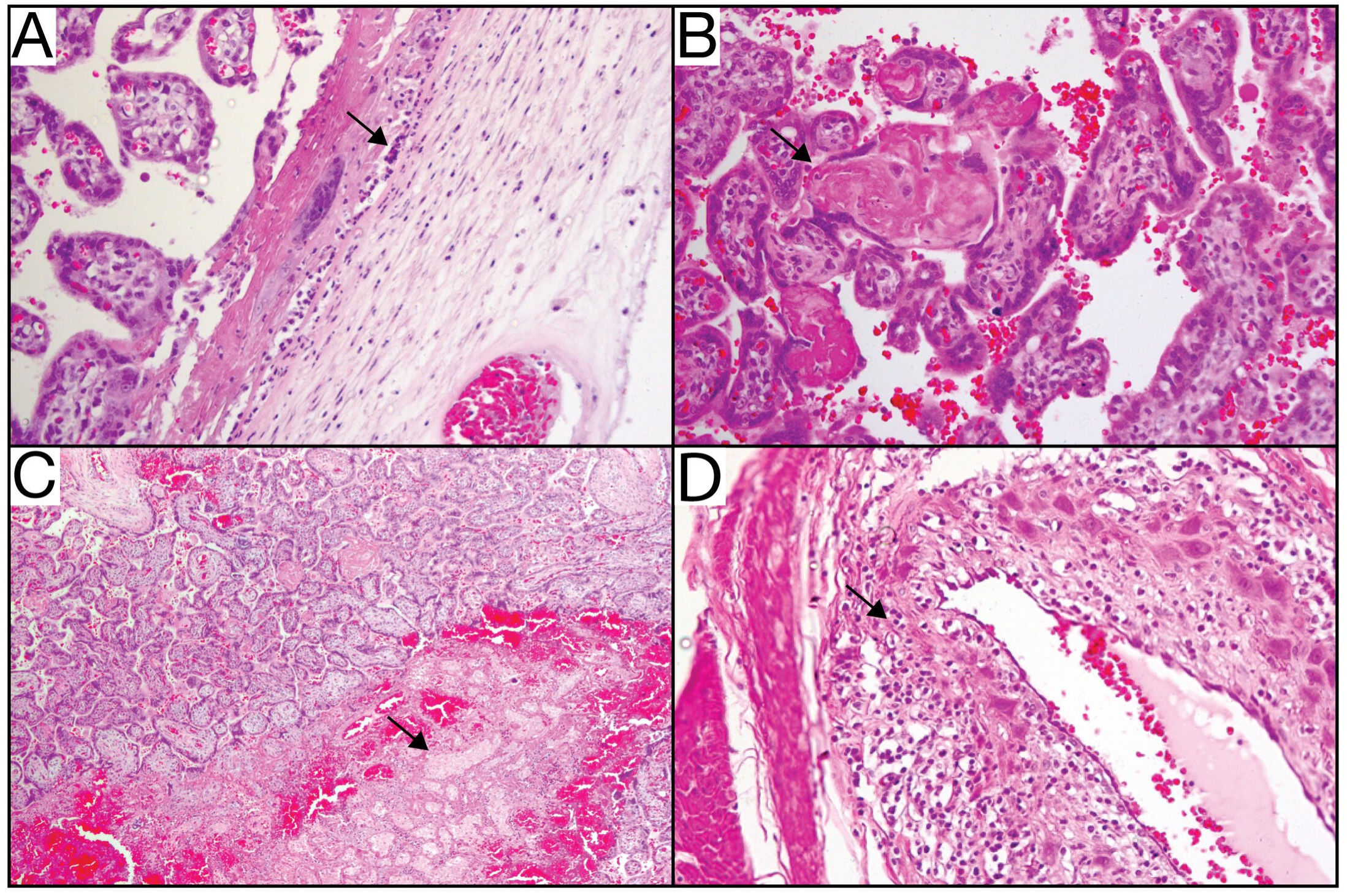
Fetal growth measured by ultrasonography. (A) Head circumference (HC), biparietal diameter (BPD), and femur length (FL) were measured in weekly ultrasounds. All measurements are depicted as millimeters (mm). The solid grey lines were derived from reference ranges from Tarantal et al. 2005 to show the mean (black lines) and one, two, and three standard deviations from the mean (grey lines). The HC, BPD, and FL were then plotted along these reference ranges to observe any deviations from the mean. Representative images of the HC, BPD, and FL ultrasounds are located to the right of the respective graph. (B) The pGA is plotted against the aGA (based on gestational age estimated from breeding and menstrual history). The pGA is shown separately for each measurement: BPD (triangle), HC (square), and FL (circle).

We also closely observed placental and fetal health by ultrasonography. No significant placental lesions were identified until 35 dpi (gd 81) when ultrasonography identified a possible area of placental abruption and a retroplacental clot along the edge of the placenta over the cervix, which was resolving by 42 dpi (gd 88). No fetal abnormalities were noted at either time point, and the fetus did not demonstrate any persistent tachycardia or bradycardia. Because of these small placental lesions, daily heart rate monitoring was initiated and remained within a normal range until 49 dpi (gd 95) when no fetal heartbeat was detected.

The dam underwent euthanasia and necropsy for a comprehensive collection of both maternal and fetal tissues. During the necropsy, the cervix was noted to be closed and no debris was noted in the vaginal vault. The amniotic sac contained significant amounts of adherent purulent matter, and purulent fibrinous material covered the decidua and fetus (Figure 6). The fetus showed advanced tissue autolysis, including severe autolysis of the fetal brain (not shown). Bacterial culture obtained by swab of the fibrinopurulent amniotic fluid at the time of necropsy demonstrated *Staphylococcus epidermidis.* We also identified clusters of gram positive cocci in the fetal esophageal lumen (Supplementary Figure 1). *S. epidermidis* is part of the vaginal flora in rhesus macaques (61). No additional samples were obtained for culture or bacterial 16s rDNA-PCR.

**Figure 6.**
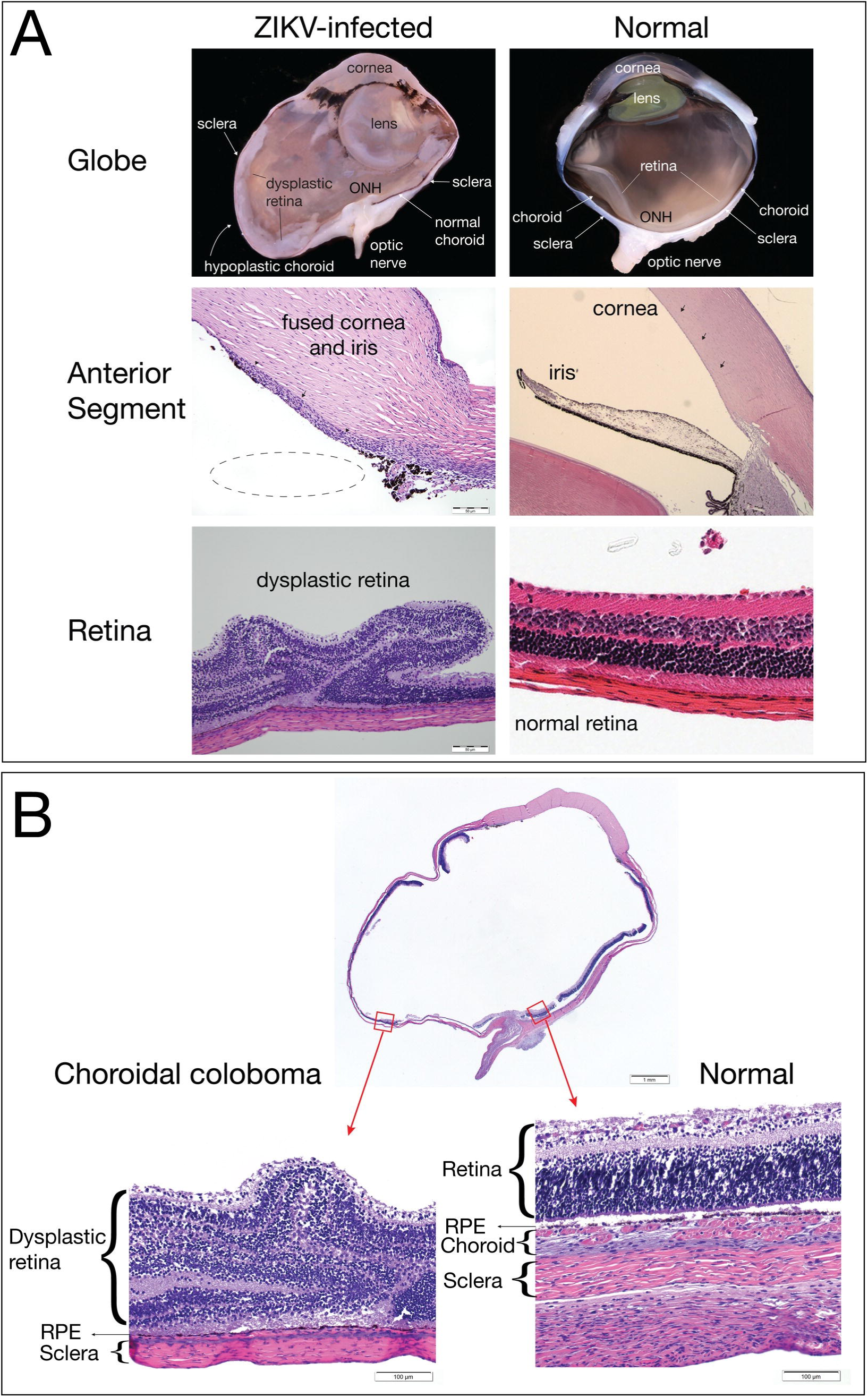
Maternal and fetal necropsy images. (A) The uterus was removed in entirety from the abdominal cavity of the dam using sterile instruments and a syringe was used to aspirate the purulent fluid from inside the uterine cavity. (B) The fetus was removed from the uterus and was covered in thick fibrinous material. (C) and (D) Placental discs 1 and 2 were covered in the same thick fibrinous maternal that covered the fetus.

### Fetal and maternal vRNA tissue distribution

At necropsy, a range of fetal and maternal tissues were processed for qRT-PCR to determine ZIKV RNA burden. ZIKV RNA was widely distributed within fetal tissues, maternal lymphoid structures, and the reproductive tract: 33 fetal, maternal and maternal-fetal interface tissues were positive for vRNA (Table 1 vRNA); of these, 27 were fetal tissues. Amniotic fluid collected during necropsy also contained vRNA (Figure 2) but did not contain replicating virus as assessed via plaque assay (not shown). The highest viral loads were detected in fetal colon and fetal lung tissue. Most organs of the fetal digestive system had detectable vRNA: stomach, jejunum, and colon. The presence of vRNA in fetal ocular structures and cerebellum indicates a central nervous system infection. vRNA was detected in four maternal lymph nodes and the spleen, indicating that ZIKV RNA was still present in the maternal immune system structures at 49 dpi, despite the absence of detectable maternal viremia. 127 maternal biopsies and fetal tissues were assayed for vRNA. All the tissues positive for ZIKV RNA are listed in this table. The maternal and fetal tissues which were vRNA negative are listed in Supplementary Table 1.

**Table 1:**
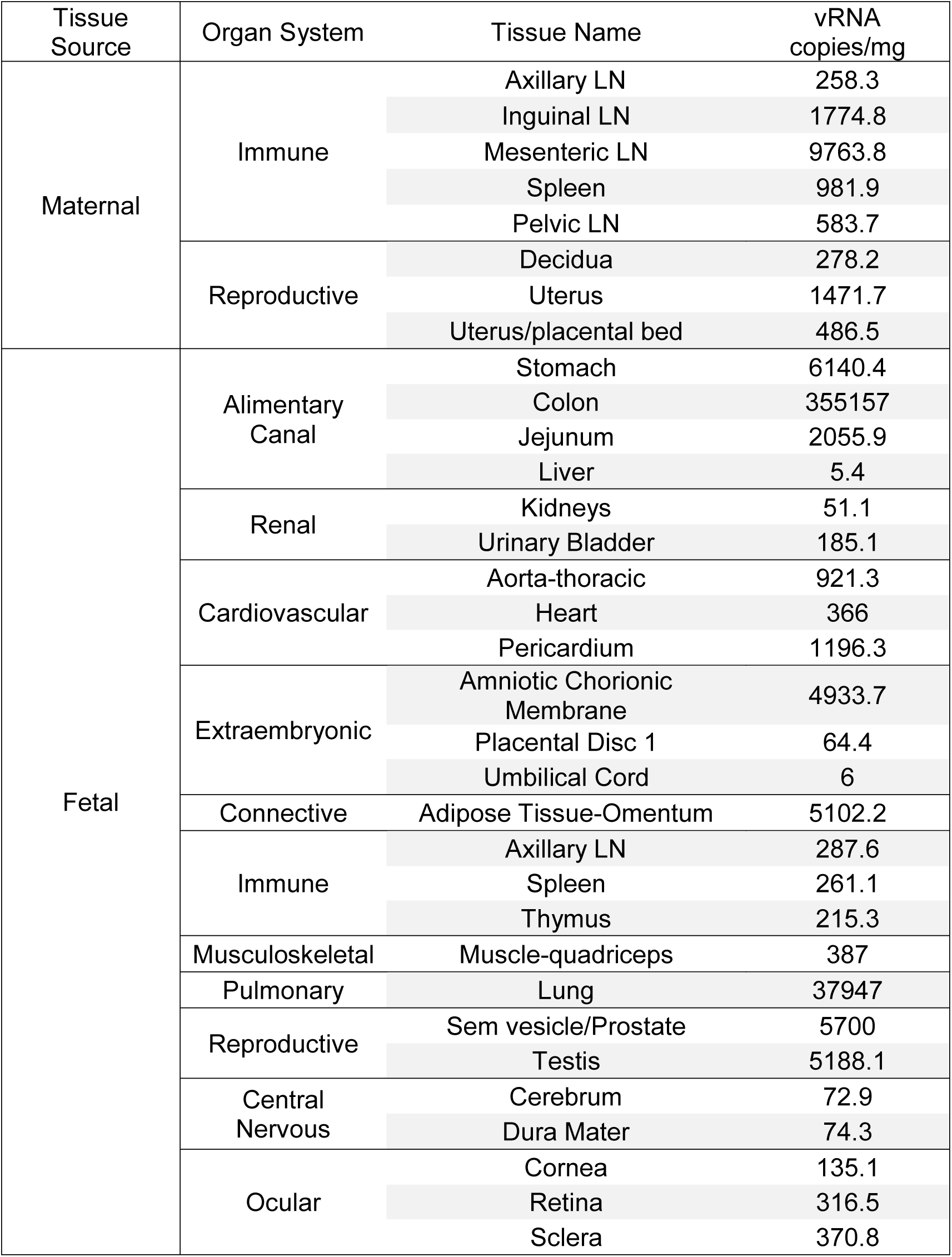
Tissues with detectable ZIKV RNA from mother and fetus

| Tissue Source | Organ System | Tissue Name | vRNA copies/mg |
| --- | --- | --- | --- |
| Maternal | Immune | Axillary LN | 258.3 |
|  |  | Inguinal LN | 1774.8 |
|  |  | Mesenteric LN | 9763.8 |
|  |  | Spleen | 981.9 |
|  |  | Pelvic LN | 583.7 |
|  | Reproductive | Decidua | 278.2 |
|  |  | Uterus | 1471.7 |
|  |  | Uterus/placental bed | 486.5 |
| Fetal | Alimentary Canal | Stomach | 6140.4 |
|  |  | Colon | 355157 |
|  |  | Jejunum | 2055.9 |
|  |  | Liver | 5.4 |
|  | Renal | Kidneys | 51.1 |
|  |  | Urinary Bladder | 185.1 |
|  | Cardiovascular | Aorta-thoracic | 921.3 |
|  |  | Heart | 366 |
|  |  | Pericardium | 1196.3 |
|  | Extraembryonic | Amniotic Chorionic Membrane | 4933.7 |
|  |  | Placental Disc 1 | 64.4 |
|  |  | Umbilical Cord | 6 |
|  | Connective | Adipose Tissue-Omentum | 5102.2 |
|  | Immune | Axillary LN | 287.6 |
|  |  | Spleen | 261.1 |
|  |  | Thymus | 215.3 |
|  | Musculoskeletal | Muscle-quadriceps | 387 |
|  | Pulmonary | Lung | 37947 |
|  | Reproductive | Sem vesicle/Prostate | 5700 |
|  |  | Testis | 5188.1 |
|  | Central Nervous | Cerebrum | 72.9 |
|  |  | Dura Mater | 74.3 |
|  | Ocular | Cornea | 135.1 |
|  |  | Retina | 316.5 |
|  |  | Sclera | 370.8 |

### Uteroplacental histopathology

Maternal-fetal interface tissues were evaluated for histological evidence of infection and lesions. There was clear evidence of both acute chorioamnionitis consistent with bacterial infection (Figure 7A), and features of relative placental insufficiency. There is no acute or chronic villitis, but the villi do show increased perivillous fibrin deposition (Figure 7B), and there are multiple remote infarctions (Figure 7C), which is a finding consistent with insufficiency. Radial arteries in the myometrium showed a pronounced leukocytoclastic vasculitis defined as an infiltrative mixture of lymphocytes, eosinophils, and plasma cells into the smooth muscle wall of these vessels (Figure 7D). The leukocytoclastic vasculitis seen around the radial arteries is usually related to hypersensitivity reactions or viral infections, and is not a consequence of bacterial infection. The decidua, placenta, placental bed and amniotic/chorionic membranes also showed significant pathology (Supplementary Table 2).

**Table 2:**
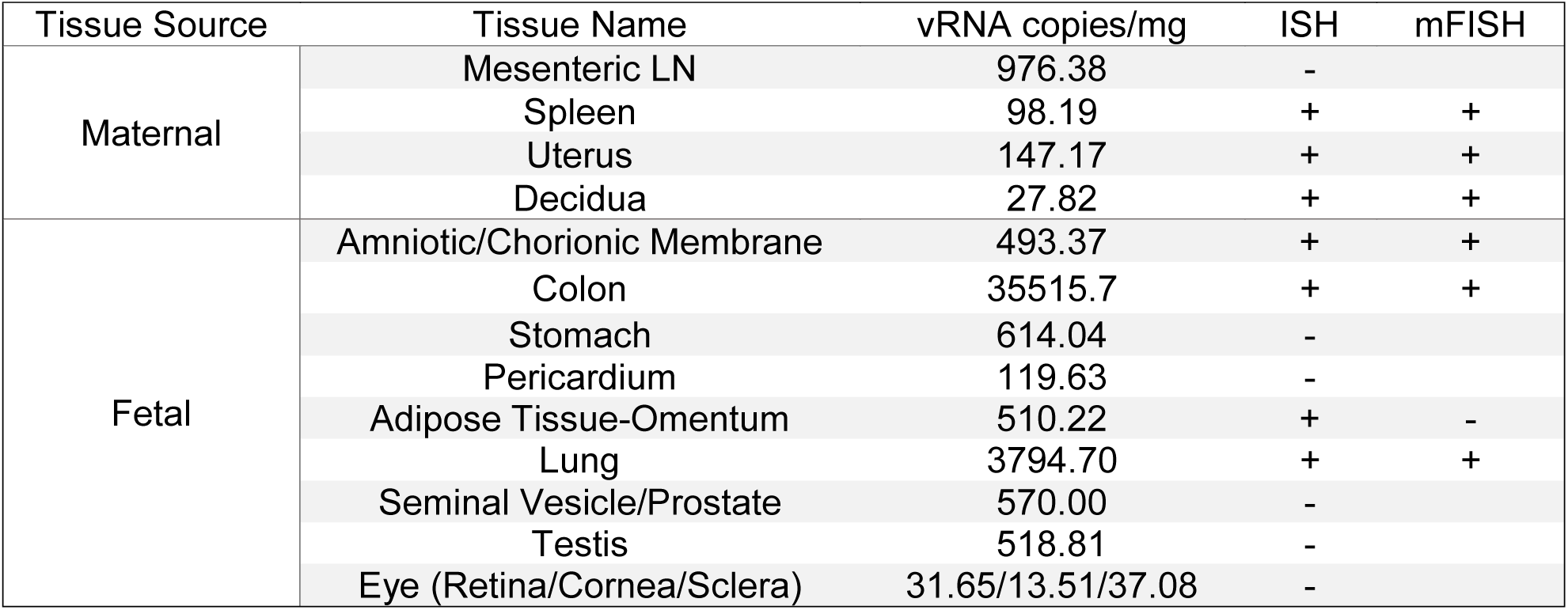
Tissue vRNA burden, ISH and mFISH results.

**Figure 7.**
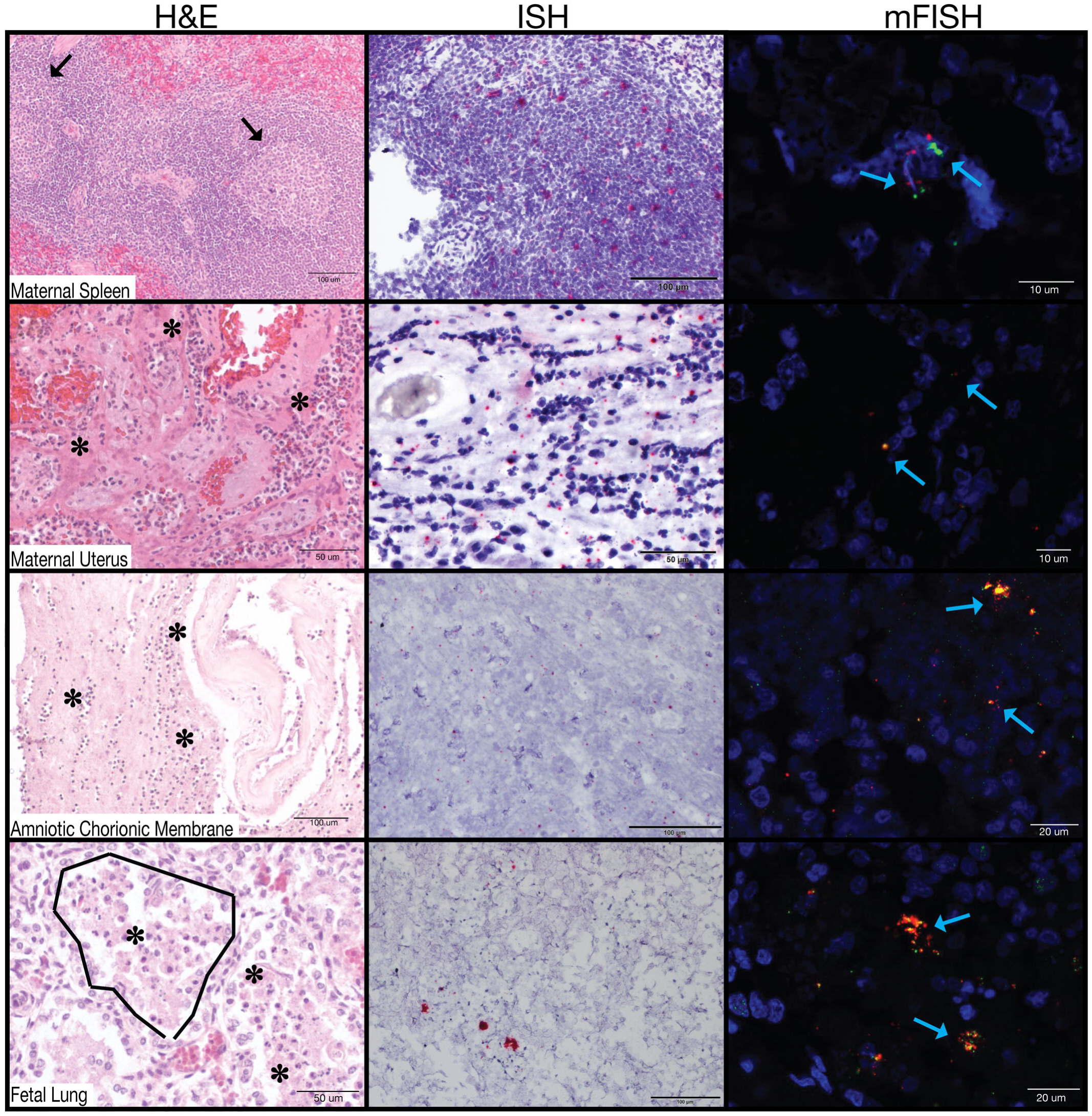
Uteroplacental histopathology. (A) Maternal neutrophils invading chorionic plate (arrow) is diagnostic of acute chorioamnionitis. (B) Villi show increased perivillous fibrin deposition (arrow) and there are multiple remote infarctions (arrow, C). (D) Radial arteries in the myometrium show a pronounced leukocytoclastic vasculitis (arrow) defined as an infiltrative mixture of lymphocytes, eosinophils, and plasma cells into the smooth muscle wall of these vessels.

### Fetal ocular histopathology

In our previous study (49), 2/2 macaque fetuses from first trimester ZIKV infection had suppurative inflammation in the ocular tissues at term (retina, choroid, optic nerve). In the current study, ocular tissues were therefore carefully evaluated by qRT-PCR and histology. One fetal eye was dissected for vRNA detection by qRT-PCR and the contralateral eye was fixed and processed for histological analysis. ZIKV RNA was detected by qRT-PCR in the retina, choroid, and lens at low levels (TABLE 1 vRNA). At the time of demise the fetal eyelids were still fused, suggesting that vRNA present in the eye was not due to passage of the virus from the amniotic fluid directly across the cornea or sclera.

In the fixed and processed globe, a chorioretinal coloboma affecting the ventral aspect of the globe was revealed, and was characterized by extensive areas of choroidal and scleral thinning with a central area of choroidal and retinal pigmented epithelium absence and marked dysplasia of the adjacent retina (Figure 8). Additionally, the presence of fusion of the iris with the posterior corneal stroma and a seeming lack of adequate maturation of the iridocorneal angle structures suggested the presence of anterior segment dysgenesis. It is necessary to acknowledge that the histologic interpretation of the anterior segment changes in this globe was hampered by the tissue autolysis presence in the fetus. Although the chorioretinal lesions were obvious even with a mild degree of autolysis, the autolytic changes impacted our ability to analyze the delicate structure of the developing tissues of the iridocorneal angles, making it impossible to definitively diagnose anterior segment dysgenesis. Because the retina is part of the central nervous system, the finding of retinal dysplasia indicates that the fetus had CNS abnormalities. The presence of the coloboma, dysplastic retina, and potential anterior segment dysgenesis are abnormalities that likely arose from disruption of early ocular developmental processes (62), consistent with the first trimester window of sensitivity in our earlier study (49).

**Figure 8.**
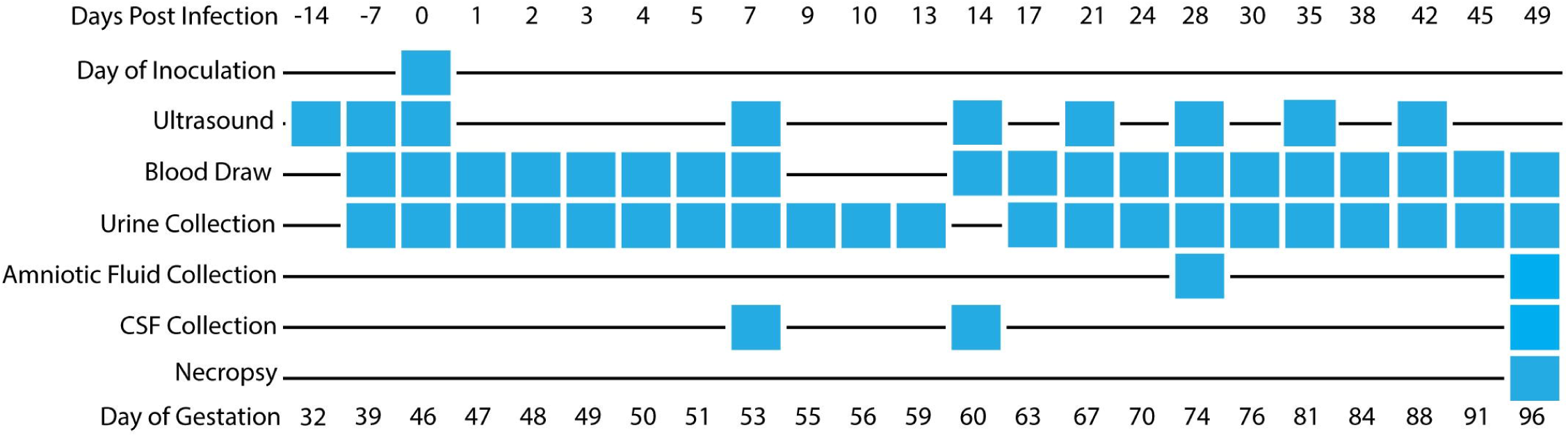
Fetal ocular pathology. (A) The left panels contain images of the ZIKV-infected eye, and the right panels show normal features from a different infant macaque for comparison. The globe of the ZIKV-infected fetus shows a hypoplastic choroid and dysplastic retina compared to the normal eye. The irregular shape of the eye in the ZIKV-infected globe is a processing artifact. The anterior segment image of the ZIKV-infected fetus shows that the iris is fused to the posterior cornea (black arrow heads), suggesting anterior segment dysgenesis; the dotted line shows where the iris would be in normal ocular development. The ZIKV-infected eye presents marked retinal dysplasia, characterized by retinal folding and loss of normal retinal organization when compared with the normal retina in the control image on the right. (B) A choroidal coloboma was identified on the ventral aspect of the globe (left image); the choroid had normal development on the dorsal aspect of the same globe (right image). The retina, retinal pigment epithelium (RPE), choroid (if present), and sclera are labeled with the left image demonstrating an absence of choroid.

### Tissue pathology and detection of vRNA in maternal and fetal tissues

Histologic lesions were noted in the fetal tissues that were potentially exposed to virus in the amniotic fluid, specifically the respiratory and gastrointestinal systems. There were significant lesions in the lungs, mesenteric lymph node, placenta, chorioamniotic membranes, decidua, maternal uterus, and maternal spleen (Supplementary Table 2). Consistent with the bacterial growth of *S. epidermidis* from amniotic fluid, gram positive cocci were observed within the lumen of the esophagus (Supplementary Figure 1), although there was no associated inflammatory reaction within the epithelium or deeper tissue layers of the esophagus. The stomach and small intestine had mucosal autolysis with no discernible histologic lesions. The lumen of the colon had granular basophilic material consistent with nuclear debris.

The fetal lungs had notable pathology. The pulmonary alveoli had fibrin, cellular debris, edema, occasional squamous cells, and neutrophilic infiltration (alveolitis). There were multiple areas of alveoli with type II pneumocyte hyperplasia, with multifocal expansion of the alveolar septa with fibrin. The trachea, primary and secondary bronchi, had relatively intact respiratory epithelium.

### ZIKV histological analyses

ZIKV RNA localization was evaluated by ISH and mFISH on selected tissues with high vRNA burden as determined by qRT-PCR. Figure 9 presents photomicrographs from near sections of the same spleen, fetal membranes, and fetal lung specimens. H&E staining is presented to demonstrate tissue organization and pathology; ISH to confirm the presence of ZIKV genome within cells, and mFISH for both negative and positive strand ZIKV RNAs to detect the dsRNA of replicative intermediates. Supplementary Figure 2 also presents representative images of positive and negative strand RNAs for the tissues displayed; the merged figure colocalizes both positive and negative sense RNA strands, indicating active ZIKV replication in these tissues.

**Figure 9.**
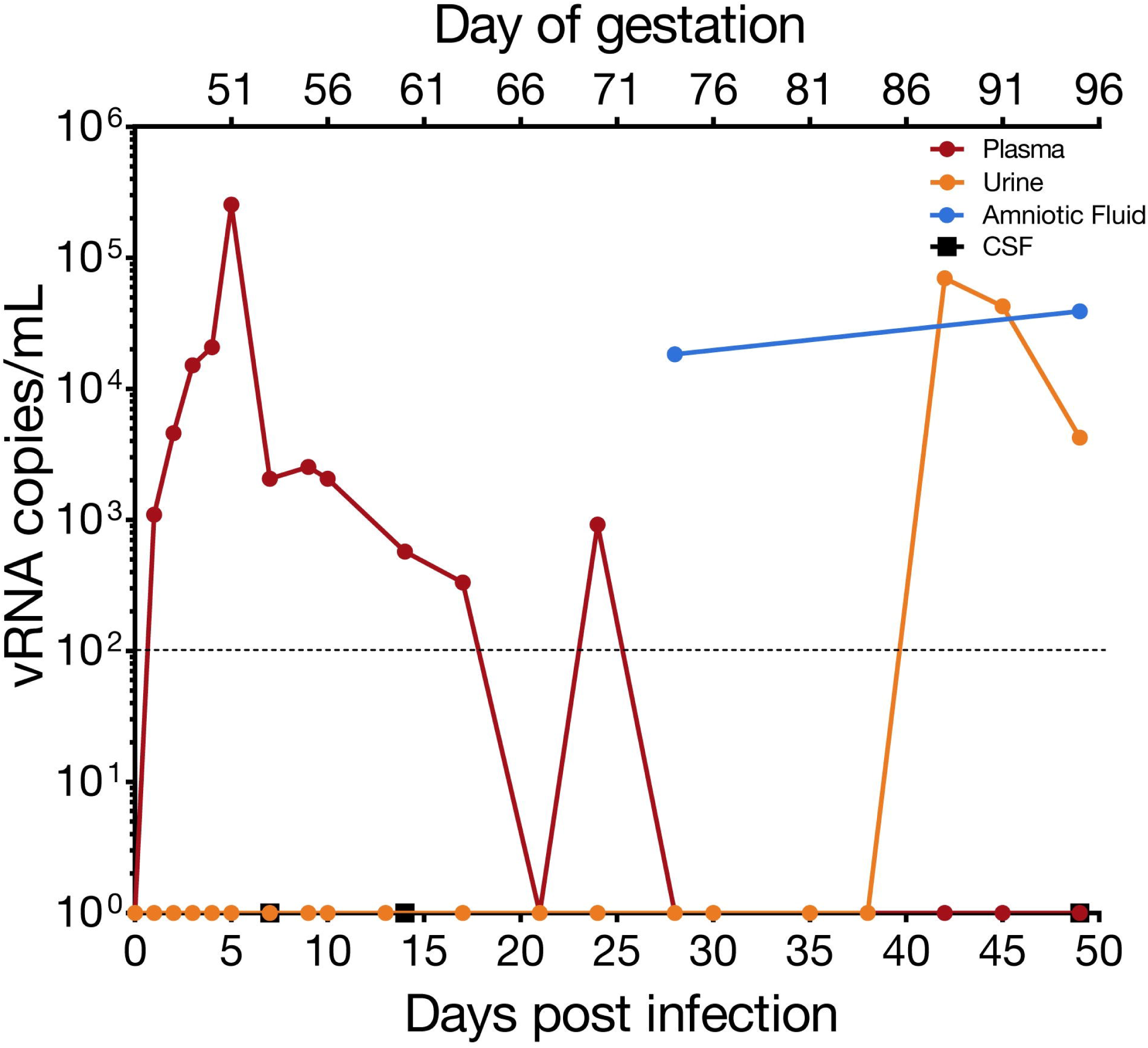
Tissue histology and viral localization of maternal spleen, maternal uterus, amniotic/chorionic membrane, and fetal lung. Each tissue was stained with H&E, ISH, and mFISH. ISH shows localization of ZIKV vRNA. mFISH shows replicative intermediates by staining the negative sense RNA strands green and positive sense RNA strands red. Colocalization (yellow) demonstrates dsRNA intermediates. Black arrows denote a germinal center. Asterisks indicate neutrophils. Blue arrows highlight green, red, or yellow fluorescence.

Tissues with a detectable vRNA burden were evaluated by ISH and mFISH. ISH detects positive sense vRNA; mFISH detects ZIKV replicative intermediates (negative and positive sense vRNA). “+” indicates detectable signal in these tissues sections, “-“ indicates no signal. Tissues with no detectable ISH signal were not further evaluated by mFISH.

## Discussion

In this report of an adverse pregnancy outcome following ZIKV infection in a rhesus macaque, we describe fetal demise following suspected PPROM, fetal and maternal ZIKV burden, and significant ocular pathology in the fetus. ZIKV RNA was widely distributed throughout fetal tissues at necropsy, including in the cerebellum and ocular tissues. ZIKV vRNA was also identified in maternal lymph nodes and maternal spleen at the time of necropsy (49 dpi). Replication competent virus was identified by ISH for the presence of negative and positive strand RNA in fetal and maternal tissues. Abnormal histology was characterized in multiple fetal tissues including alveolitis and pneumocyte hyperplasia in fetal lung tissue, and severe ocular abnormalities. Both fetal ocular pathology and fetal demise have been described in human reports of ZIKV infection and demonstrate parallels between human and NHP CZS.

### Fetal demise

This is the first report of rupture of membranes and fetal demise in an NHP model of congenital ZIKV infection. We presume that maternal membranes ruptured around 42 dpi (although some amniotic fluid may have been present at 28 dpi) because we detected amniotic fluid markers in the urine at this time point, and identified high vRNA burden in this urine sample, despite absence of detectable maternal viremia at this time. A week after detection of ZIKV RNA in the pan-collected urine/amniotic fluid mixture, abdominal ultrasound evaluation found no fetal heartbeat and the fetus and dam were submitted for necropsy. There was no chronic villitis, which would be expected for viral induced changes. However, sections of the decidua and myometrium revealed a pronounced leukocytoclastic vasculitis involving the smooth muscle walls of the radial and spiral arteries. This is significant because this type of vasculitis is not expected in cases of bacterial infection, but do occur as a response to viral infections associated with cutaneous vasculitis (hypersensitivity vasculitis) (63).

Although there was fibrinopurulent material surrounding the fetus in the uterine cavity and gram positive cocci in the esophagus, multiple sections of placenta and all other fetal tissues had no histologic evidence of bacterial colonization. The growth of *S. epidermidis* from aspirated amniotic fluid was minimal and contamination at the time of collection is a possibility. Neutrophilic infiltration, such as that seen in the chorionic plate, is consistent with bacterial infection, but intraamniotic neutrophilic inflammation can also be sterile (64). Sterile neutrophilic inflammation has been reported previously in experimental infection in animals models with this strain of ZIKV, including mice which demonstrated neutrophil infiltration of the skeletal muscle and hippocampus (65) and male rhesus macaques which demonstrated interstitial neutrophilic prostatitis (47). Therefore, while this clinical presentation is consistent with an ascending bacterial intraamniotic infection, further studies will be able to provide clarification of the histopathologic outcomes with macaque pregnancies.

Closely associated with this fetal demise is the occurrence of PPROM. Although it is not possible to determine if the amniotic membranes ruptured because of ZIKV infection, the finding of PPROM followed by fetal demise also occurs during human prenatal ZIKV infection (24). It could be hypothesized that the amniocentesis at 28 dpi contributed to the possible intrauterine bacterial infection; however, the long duration of time separating these events, and the typical rapidity of preterm labor in the rhesus macaque with experimental bacterial infection of the amniotic fluid makes this unlikely (66, 67). Additional studies of ZIKV infection during NHP pregnancy are needed to determine if there is an association between congenital ZIKV infection and an increased risk for intra-amniotic infection leading to PPROM and fetal demise. One may speculate that ZIKV infection early in gestation affects pregnancy-induced T-cell changes or placental invasion involved in uterine vascular remodeling necessary for normal blood flow to the placenta. In turn, ZIKV infection may lead to abnormal remodeling and abnormal blood flow to the placenta culminating in pathologic infarctions and increased risk for relative placental insufficiency and preterm birth. This working hypothesis requires further study.

### Fetal ocular defects

Congenital ocular abnormalities are strongly associated with human prenatal ZIKV infection, as demonstrated by the high frequency (up to 55%) of ocular disease in human infants with first trimester prenatal infections (12). There is growing evidence that structures of the fetal visual system are a significant target for ZIKV in human pregnancy. The fetal eye evaluated for pathology in the current study had anterior segment dysgenesis, a ventral choroidal coloboma, and retinal dysplasia. This is the first time that such severe ocular abnormalities have been reported with macaque CZS. As far as we are aware, bacterial infections are not associated with such abnormalities during development. In addition, an acute intrauterine bacterial infection would not have impacted eye development, since the ocular structure damage described would likely have occurred from the disruption of normal developmental processes which occur earlier in pregnancy. Anterior segment dysgenesis refers to a spectrum of developmental anomalies resulting from abnormalities of neural crest migration and differentiation during fetal development (68). In humans, anterior segment dysgenesis is present in rare syndromes (69), and although the rate of anterior segment dysgenesis and related syndromes is unknown in rhesus macaques, it would be unlikely to appear in pregnancy.

An ocular coloboma is a congenital lesion associated with a failure in the closure of the embryonic (ocular) fissure causing defects of one or more ocular structures (i.e., the eyelids, lens, cornea, iris, ciliary body, zonules, choroid, retina and optic nerve). The defect is essentially a bare sclera with the overlying retinal pigmented epithelium, retina or choroid missing (70). It may be sporadic or inherited and, in some cases, is associated with systemic disorders (70). Choroidal colobomas in humans can be also associated with the presence of retinal dysplasia (71, 72), which was noted in the current case. Although there are multiple genetic mutations associated with colobomatous defects in humans (70), there is only one case report of a macaque with coloboma and no genetic evaluations were pursued in that report (73). We did not pursue a genetic evaluation because it seems unlikely that a rare genetic defect would occur in one of the fetuses with congenital ZIKV infection. The defects in the eye affected the posterior and ventral aspect of the globe, which is common, since the ocular fissure is embryologically located in the ventro-nasal quadrant of the eye. It also mainly affected the choroid, thereby classifying it as a choroidal coloboma. In our previous study, we identified optic nerve gliosis in the two-first trimester infections (49), but did not identify other significant ocular pathology. CZS represents a continuum of disease from mild to severe and the macaque model highlights this by capturing the wide disease spectrum. It is also important to note that the current study was conducted with a virus stock prepared from an isolate obtained from a person infected in Puerto Rico, whereas our previous study (49) was conducted with a virus stock prepared from a French Polynesian isolate. Our results may indicate that closely related viruses can cause different outcomes in pregnant macaques, however further studies will be needed to understand whether specific genetic determinants are related to these outcomes. Although ZIKV causes ocular disease in the adult murine model (74), no ocular anomalies to this extent have yet been observed in mouse models of congenital ZIKV infection. This also underscores the important role the rhesus macaque model plays in studying ZIKV effects on pregnancy outcomes.

### Maternal and fetal tissue viral distribution

ZIKV RNA was detected throughout fetal tissues, affecting multiple organ systems (digestive, respiratory, reproductive, cardiovascular, immune, and nervous), and replication competent virus was identified in fetal lung tissue 49 dpi via negative and positive strand RNA ISH. Remarkably, ZIKV RNA was also detected in maternal lymph nodes at 49 dpi and replication competent virus was identified in the lymph node tested. The presence of vRNA does not imply that the virus is replicating or may be transmissible. However, the detection of negative strand RNA by ISH and its colocalization with positive strand vRNA is confirmation of replication competent virus, and the finding of infectious virus in fetal and maternal tissues 49 dpi could have important implications for transmission. The fact that replication competent ZIKV is still present in an adult lymphoid-associated tissue at 49 dpi is critical to understanding the risk involved with organ transplantation, although with the caveat that viral persistence may be longer in pregnancy. Viral persistence is not explained by a lack of maternal humoral immune response since the dam developed neutralizing antibodies at concentrations similar to our previous NHP studies of ZIKV infections (50, 55).

We do not know to what extent ZIKV infection of the fetus directly contributed to fetal demise, since this case is complicated by PPROM with acute chorioamnionitis. Extended exposure of the fetus to ZIKV is most likely responsible for the ocular pathology observed, and there is no literature of which we are aware which suggests that bacterial infection results in ocular malformations. The substantial viral burden in the fetal membranes also supports the hypothesis that ZIKV contributed to PPROM. The detection of replicating ZIKV intermediates in membranes and fetal tissues at the time of fetal demise also suggests that active ZIKV infection was ongoing up until fetal demise.

In summary, we describe a case of congenital ZIKV infection with severe ocular and uteroplacental pathology complicated by fetal demise following apparent PPROM. The fetal ocular pathology recapitulates defects seen in human CZS. This is the first report of an adverse pregnancy outcome and fetal pathology in an NHP infected with ZIKV strain PRVABC59, and thus supports the importance of the macaque model for not only defining the risk ZIKV poses for pregnant women and their fetuses in the Americas, but also for defining the precise pathways by which ZIKV accesses the fetal compartment, and for testing strategies to intervene in vertical transmission.

